# Traction Force Microscopy by Deep Learning

**DOI:** 10.1101/2020.05.20.107128

**Authors:** Y.L. Wang, Y.-C. Lin

## Abstract

Cells interact mechanically with their surrounding by exerting forces and sensing forces or force-induced displacements. Traction force microscopy (TFM), purported to map cell-generated forces or stresses, represents an important tool that has powered the rapid advances in mechanobiology. However, to solve the ill-posted mathematical problem, its implementation has involved regularization and the associated compromises in accuracy and resolution. Here we applied neural network-based deep learning as a novel approach for TFM. We modified a network for processing images to process vector fields of stress and strain. Furthermore, we adapted a mathematical model for cell migration to generate large sets of simulated stresses and strains for training the network. We found that deep learning-based TFM yielded results qualitatively similar to those from conventional methods but at a higher accuracy and resolution. The speed and performance of deep learning TFM make it an appealing alternative to conventional methods for characterizing mechanical interactions between cells and the environment.

**Statement of Significance:** Traction Force Microscopy has served as a fundamental driving force for mechanobiology. However, its nature as an ill-posed inverse problem has posed serious challenges for conventional mathematical approaches. The present study, facilitated by large sets of simulated stresses and strains, describes a novel approach using deep learning for the calculation of traction stress distribution. By adapting the UNet neural network for handling vector fields, we show that deep learning is able to minimize much of the limitations of conventional approaches to generate results with speed, accuracy, and resolution.

## Introduction

Mechanobiology examines the input and output of mechanical forces by cells. Contractile forces generated by the actin-myosin cytoskeleton are transmitted to the exterior environment via associated trans-membrane proteins such as integrins (1). Referred to as traction forces, these forces are detected predominantly near cell periphery or front in an orientation towards cell center (2). Active forces are located at nascent or maturing focal adhesions near the front while resistive forces are located at mature focal adhesions near the rear (3, 4).

In addition to propelling cell migration, traction forces are believed to perform important functions such as organizing the extracellular matrix (5), probing mechanical properties of the environment (6), and sensing the state of the cell itself such as shape and size (7). Adhesive cells are also keenly sensitive to mechanical forces transmitted via the surrounding matrix (8), fluid shear (9), or cell-cell contact (10). These mechanical signals elicit many profound responses to affect cell migration (11), growth (12), and differentiation (13).

Methods for mapping cellular traction forces, referred to in general as traction force microscopy (TFM; 2, 14, 15, 16), represent a fundamental tool for mechanobiology. TFM is commonly performed by culturing cells on a substrate of elastic material such as polyacrylamide embedded with particle markers for mapping force-induced strains, which are then used for calculating the distribution of stresses. However, while the calculation of strain from stress is straightforward, the inverse calculation is known mathematically as an ill-posed problem, where a unique solution may not exist and the results are prone to artifacts from the noise in strain measurement (17). To mitigate these problems, most conventional approaches include a regularization term in the fitting function, which decreases the noise and complexity of the solution while also limiting the accuracy and resolution (2, 17). An additional challenge is the difficulty to define the relative weight, represented by a parameter λ, between regularization and accuracy (18).

Neural network-based deep learning has been deployed as a powerful method for solving ill-posed problems (19). It involves the optimization of a cascade of convolutional operations, with the goal of transforming the input (e.g. the distribution of strains) into the targeted solution (e.g. the distribution of stresses; 20). As an approach fundamentally different from conventional methods, it can avoid the use of regularization and associated compromises.

To deploy deep learning for TFM, we have adapted a neural network for processing images to process the vector fields of stresses and strains. In addition, we generated a large set of strain and ground truth stress fields by computer simulation for training the network. By comparing the performance of the resulting neural network and a conventional implementation of TFM, we show that deep learning based TFM are capable of mapping traction stress at a high speed, accuracy, and resolution.

## Materials and Methods

### Generation of Simulated Data for Training and Testing the Neural Network

Simulated stress fields were generated by modifying the model of cell migration as described by Satulovsky et al. (21). The model implemented the Local Excitation-Global Inhibition (LEGI) mechanism for controlling the protrusion and retraction activities (22), using two numbers to represent the respective signals at each location around the cell periphery. We then assumed that stress at a given location was proportional to the absolute value of the difference between these signals, and pointing at a direction toward the center of the cell. Stress vectors were placed at random distances from the cell periphery following Poisson distribution. Only simulated cells with net stresses balanced to within 10% were selected for training and testing. Some of the point stresses were blurred into patches by convolving with a 3×3 Gaussian kernel.

Substrate strains were calculated from traction stresses as described in Dembo and Wang (2) based on a Poisson ratio of 0.45, using 2D convolution with Green’s tensors followed by matrix addition to obtain net strains. Green’s tensors were calculated according to Boussinesq’s equations (2), except for the central, singular element. Since the performance of deep learning TFM was to be compared against FTTC, the value of the central element was determined by trial and error such that the value of stress was recovered as closely as possible by FTTC. For testing the performance of TFM, stresses and strains of simulated cells were scaled such that the average strain within each simulated cell, measured in modeling pixels, matched that of experimental measurements. Noise of strains were created using a Gaussian random number function at a specified standard deviation and added to the strains.

### Traction Force Microscopy

FTTC was performed using an open-source plug-in for ImageJ (23). The output was exported into MATLAB for analysis. Deep learning TFM was performed with MATLAB version 2019b. UNet was constructed layer by layer, using 3D convolution for all the convolution operations (Supplemental Fig. 1). Input and output matrices were of a dimension of 104×104×2, where the first two dimensions denoted the position of modeling pixels and the third dimension represented x and y components of the vector. Training was conducted using stochastic gradient descent with momentum (SGDM) for 200 epochs with a minibatch size of 64 and L2 regularization factor of 5×10^-4^. The forward loss function was based on half mean squared error, calculated as the square of the length of the difference vector between calculated and ground truth stress vectors, summed over all the modeling pixels and averaged over the minibatch. The learning rate was initially set at 3×10^-3^ and decreased every 10 epochs by a factor of 0.7943. Using a Dell Precision E5450 laptop computer with an Intel Xeon E-2176M CPU running at 2.70 GHz, 32 GB of RAM, and an NVIDA Quadro P2000 GPU, the training process was completed in 6 hours with a steady decline of the root mean squared error to reach 3 after 200 epochs (Supplemental Fig. 2).

### Substrate Preparation and Cell Culture

Polyacrylamide (PAA) substrates were prepared by mixing 8% acrylamide (Bio-Rad, Hercules, CA), 0.1% bis-acrylamide (Bio-Rad), and 0.2 μm fluorescent beads at a dilution of 1:1000 (Molecular Probes, Carlsbad, CA). After adding a photo-initiator, lithium phenyl-2,4,6-trimethylbenzoylphosphinate (LAP; Allevi, Philadelphia, PA) at 1 mg/ml, 25μl of the mixture was pipetted onto a no. 1 coverslip pretreated with 0.3% (v/v) bind-silane in 95% ethanol (Cytiva, Marlborough, MA). A 22mm coverslip coated with activated gelatin was placed on top^7^, and the mixture was exposed to 365 nm UV for 3 min under a tabletop UV lamp with 15W tubes (Cole Parmer, Vernon Hills, IL). The top coverslip was then gently removed with a pair of fine tweezers. Young’s modulus of the gel was measured as 10.67 kPa. The substrate was sterilized with 260 nm UV for 20 min before use. NIH 3T3 cells were maintained in Dulbecco’s modified Eagle’s medium with 10% donor bovine serum in an incubator with 5% CO2 at 37°C, and plated on PAA substrate overnight for TFM measurements.

### Measurement of Substrate Strain

To measure substrate strains generated by the traction stress, isolated NIH 3T3 cells were identified and images of fluorescent beads were recorded with a Nikon Eclipse Ti microscope equipped with a dry 40x/N.A. 0.75 PlanFluor phase contrast objective lens, before and after removing the cell by adding 1 ml of 0.25% Trypsin-EDTA solution. Images were captured with a Photometrics Prime 95B CMOS camera with 1200×1200 camera pixels. At the magnification used, each camera pixel represented an area of 0.625×0.625 μm^2^. The average surface density of beads was 14.55 / 100 μm^2^.

Bead images were cropped to 728×728 camera pixels with the cell located near the center. Pairs of images before and after cell removal were analyzed with particle image velocity using an open source plug-in for ImageJ (23). Strain vectors were generated at a spacing of 7 camera pixels and exported to MATLAB, where the image was scaled down 7x to 104×104 modeling pixels. Thus, the length of each modeling pixel equals 7x camera pixels, and each modeling pixel represented an area of 1.925 x 1.925 μm^2^ at the magnification used.

For the measurement of background strains and average strains in the cellular regionl, an outline was drawn manually around the cell at a distance of 20-40 μm from the visible border. The region within the outline was referred to as the cellular region. This ensured the inclusion of regions reached by filopodia that were invisible under the optical condition. The region further away by 40 μm was referred to as the extracellular region. Strain vectors in the extracellular region were first filtered with a 7×7 2D median filter and the average vector was calculated and subtracted from all the strain vectors such that the average extracellular strain equals zero. The standard deviation of extracellular strain vectors was used as a measure of noise in strain measurement.

### Statistical Analysis

TFM was tested with 28 simulated cells and 15 experimental measurements. For simulated cells, the accuracy of TFM was assessed against the known ground truth, with normalized squared error defined as:

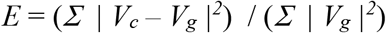

where *V_g_* was the ground truth stress vector and *V_c_* was the corresponding calculated stress vector. The difference between *V_c_* and *V_g_* was calculated using vector algebra and summed over all the modeling pixels within the cell border. Mean normalized squared error 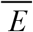, was then calculated as the average of *E* over the 28 simulated cells. Two-tail t-test for paired samples was used for determining the statistical significance of differences in mean normalized squared errors. Since ground truth stress field was not available for experimental measurements, the accuracy of TFM was assessed by comparing measured strains against strains calculated from the stress field generated by TFM, using an equation similar to that for stress vectors.

## Results

### Construction of Deep Learning Neural Network for Traction Force Microscopy

We have adapted UNet, a neural network developed for segmenting biomedical images (24), for 2D TFM. Vector field of stress or strain was represented with a 3D matrix [x, y, v], where x and y ∈ [1,104] were 2D position of the modeling pixel (to be distinguished from camera pixel) and v ∈ [1, 2] represented the two orthogonal vector components. In addition, all 2D convolution operations in UNet were replaced with 3D convolution operations. The resulting neural network for TFM contained a total of 51 layers in 3 encoding levels and 4 decoding levels, with 3 concatenation skip connections for preserving the spatial resolution (24; Supplemental Fig. 1, vertical arrows).

### Generation of Datasets for Training and Testing the Neural Network

While one may use experimental data for training the neural network for TFM, the collection of strain data from hundreds of cells will be demanding and the stress fields calculated with conventional TFM may deviate substantially from the ground truth due to regularization (17). To overcome the challenge, we have adapted a computational model for cell migration to generate simulated stress fields (21; Fig. 1a-b). Strain field was then calculated according to linear elastic theory (2; Fig. 1c). Training, conducted with 708 simulated cells for 200 epochs on a Dell Precision E5450 laptop computer, lasted for approximately 6 hours when the root mean squared error converged steadily to approximately 3 (Supplemental Fig. 2).

**Figure 1.**
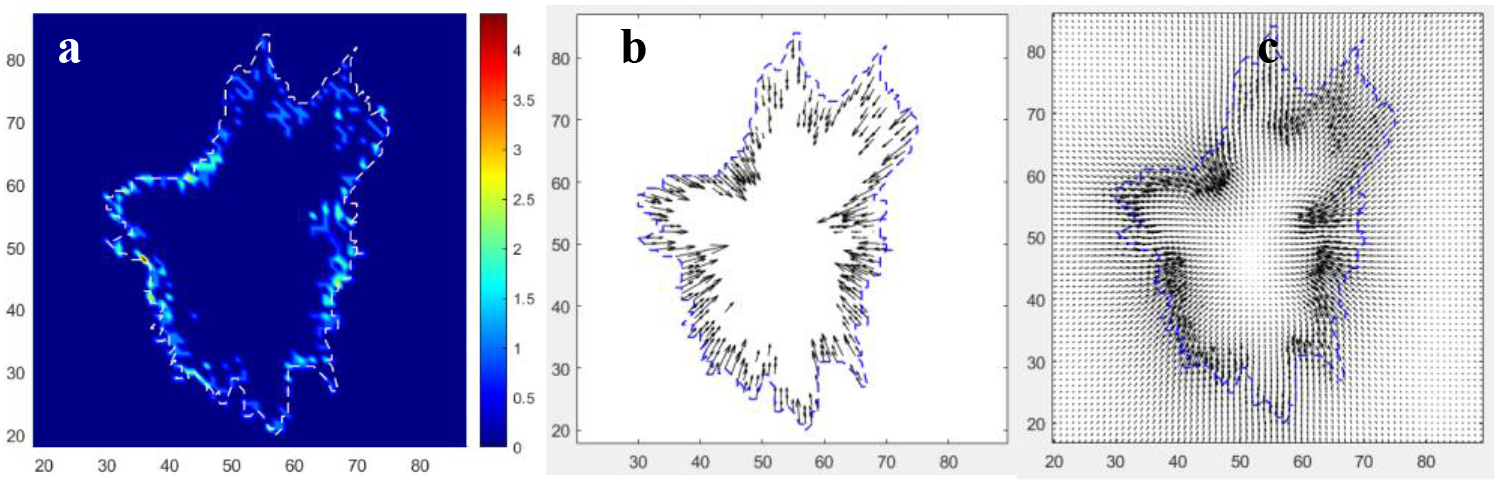
An example of simulated stress and strain fields for training the neural network. Heat map (a) and vector plot (b) show the traction stress field. Color scale is in the unit of Newtons/pixel^2^. Strains (c) are calculated assuming a Poisson ratio of 0.45 and Young’s Modulus of 10 Newtons/pixel^2^. Segmented lines indicate simulated cell border.

### Deep learning traction force microscopy for simulated data

Deep learning TFM was first tested with simulated data generated in a manner similar to training data, rescaled such that the average strain within the simulated cell border matched that of experimental measurements, in modeling pixels. The computation of each 104×104 stress field using deep learning TFM took < 1 millisecond. Heat maps of predicted stress field and ground truth showed a strong qualitative similarity (Fig. 2d and 2e). Similar heap maps were also obtained using a conventional TFM implemented as Fourier Transform Traction Cytometry (23; FTTC; Fig. 2c and 2f), although errors were more prominent with FTTC as revealed in vector plot (Fig. 2i, arrows).

**Figure 2.**
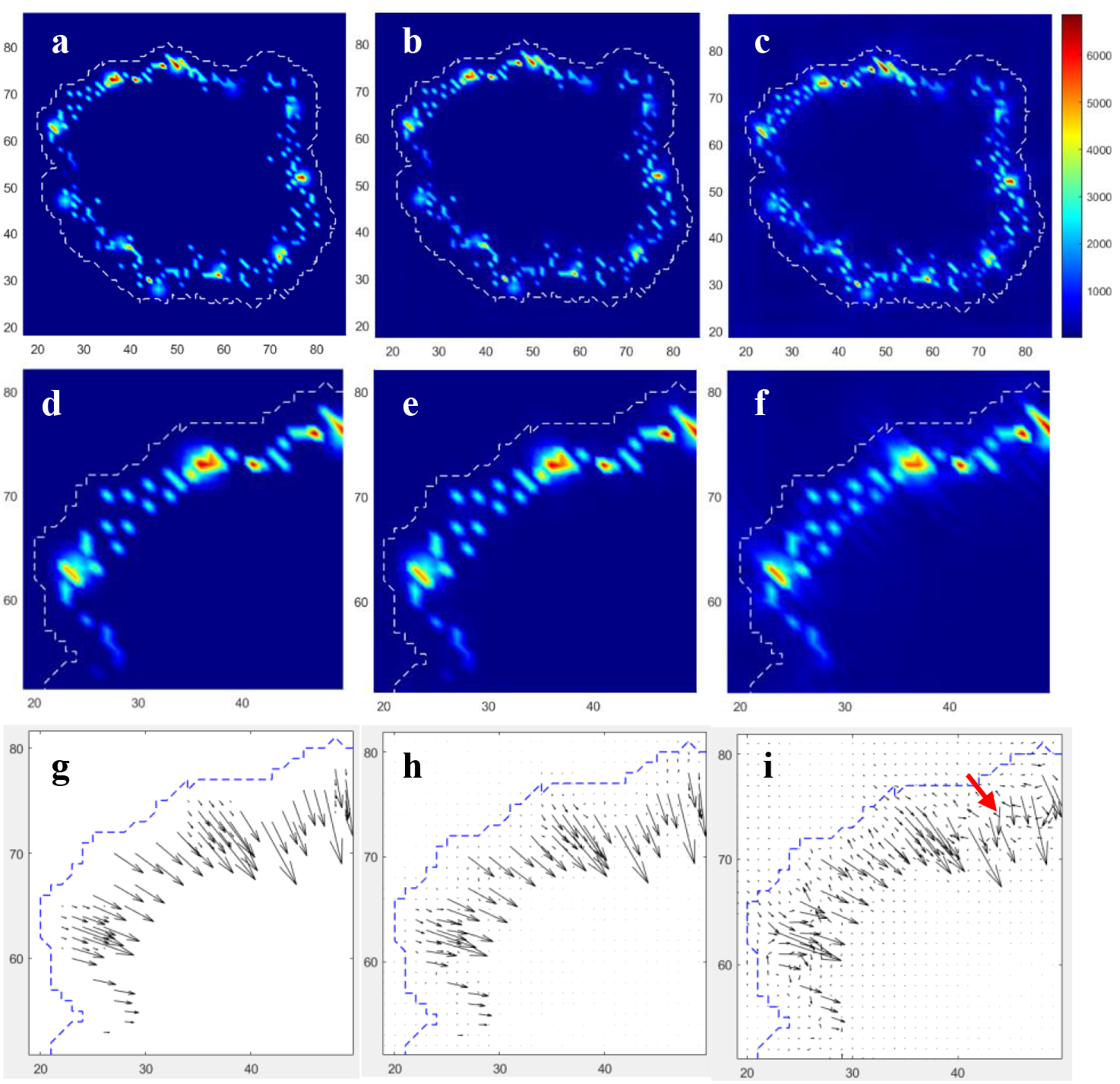
TFM of a simulated testing cell without adding noise to the strains. The resulting stress fields are shown as heat maps at a low (a-c) or high (d-f) magnification, or as vector plots (g-i). Traction fields qualitatively similar to the ground truth (a, d, g) are obtained with either deep learning (b, e, h) or FTTC (c, f, i). However, vector plot of FTTC field showed some errors (i, red arrows), and a higher background that appears as small dots or short segments (i). To mimic experimental data, each modeling pixel is assumed to represent 1.925 μm and the average length of intracellular strains is scaled to 0.4277 modeling pixels, equivalent to 0.823 μm in experimental images. Young’s Modulus of the substrate is assumed to be 10,670 Pascals. Color scale is in the unit of Pascals.

To quantify the accuracy of TFM we calculated normalized error 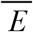, defined as the sum of squared errors for stress vectors over all the pixels divided by the sum of squared ground truth stress vectors. Without adding noise to strain vectors, 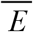 for deep learning TFM was 3.17 +/- 0.82%, while 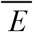 for FTTC without regularization was approximately 5 times higher at 15.1 +/- 1.99% (p≪0.0001). No improvement was observed when applying regularization to FTTC with noise-free strains (Fig. 3, left set, solid bars). The small error of deep learning TFM also suggested negligible over-fitting of the network, a common caveat of deep learning that compromises general accuracy in attempt to minimize the error of training data.

**Figure 3.**
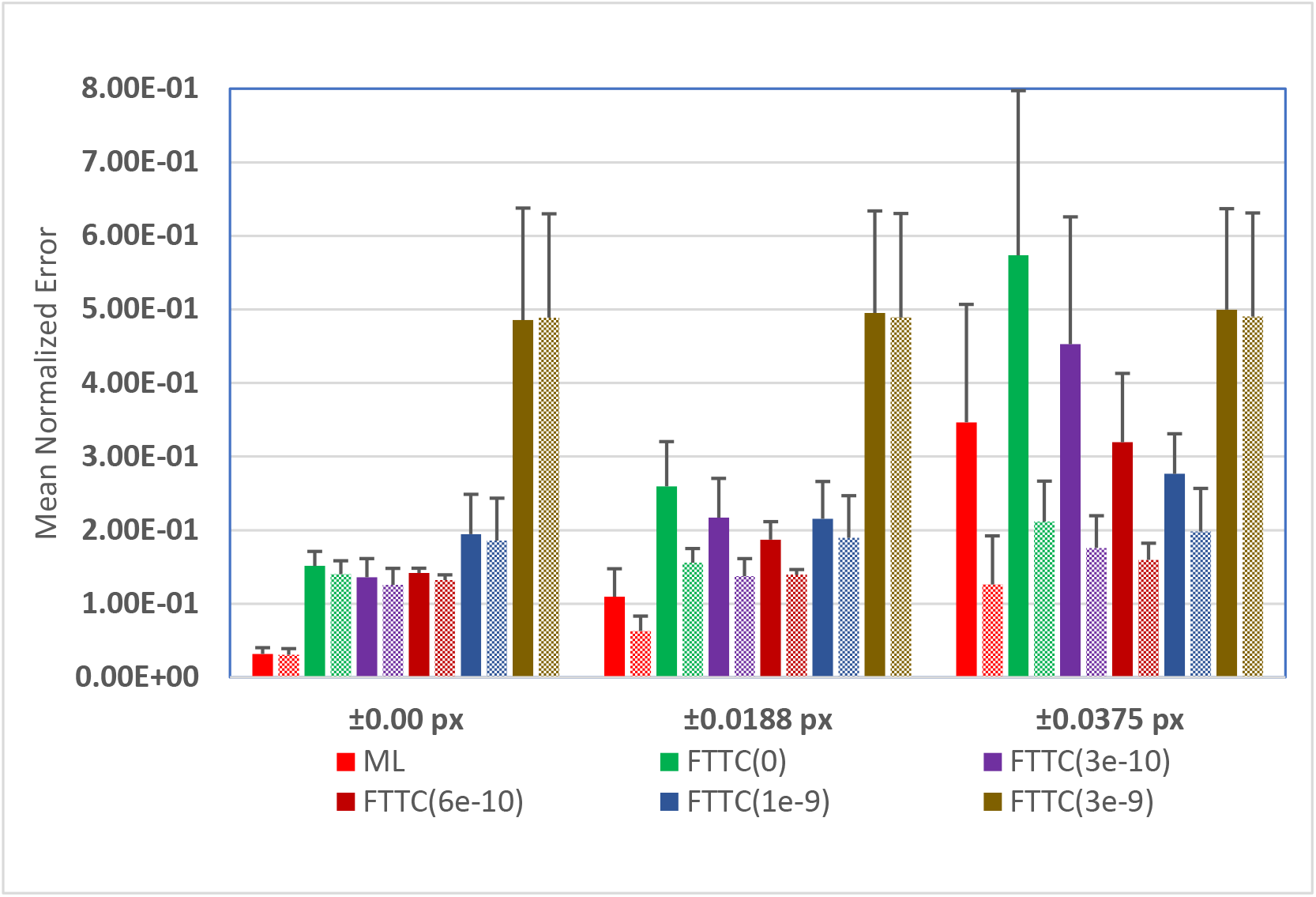
Mean normalized errors of traction stress fields as calculated with deep learning TFM (red bars) or FTTC at various levels of regularization (λ indicated in parentheses). Y values represent the mean of 28 simulated cells with no noise (left group), a noise of 0.0188 modeling pixels (equivalent to the noise in experimental measurements; middle group), or a noise of 0.0375 modeling pixels (right group) imposed on simulated strains. Dark solid bars indicate errors without applying cutoff filter; light textured bars indicate the corresponding errors after applying cutoff filter. Error bars represent standard deviations.

We have tested the effect of noise equivalent to that of experimental measurements (+/-0.0188 modeling pixels, equivalent to 0.132 camera pixels or 0.0362 μm; Fig. 3, middle set) or 2x experimental measurements (+/-0.0375 modeling pixels, equivalent to 0.263 camera pixels or 0.0722 μm; Fig. 3 right set). As expected, adding noise to strain vectors caused an increase in 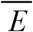 for both deep learning TFM and FTTC, while 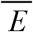 for deep learning was smaller than 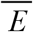 for FTTC without regularization (Fig. 3 middle and right set, solid red and green bars, p ≪ 0.0001). With regularization, 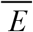 for deep learning remained smaller than that for FTTC except at 2x the experimental noise and λ = 1×10^-9^ (Fig. 3 right set, solid red and solid blue bars, p < 0.02). However, heat maps showed a loss of fine features and resolution at this level of regularization (Fig. 4a, 4c, 4d, 4f; see Fig. 2a and 2d for ground truth), while qualitative features were preserved using deep learning TFM (Fig. 4a, 4d, 4j).

**Figure 4.**
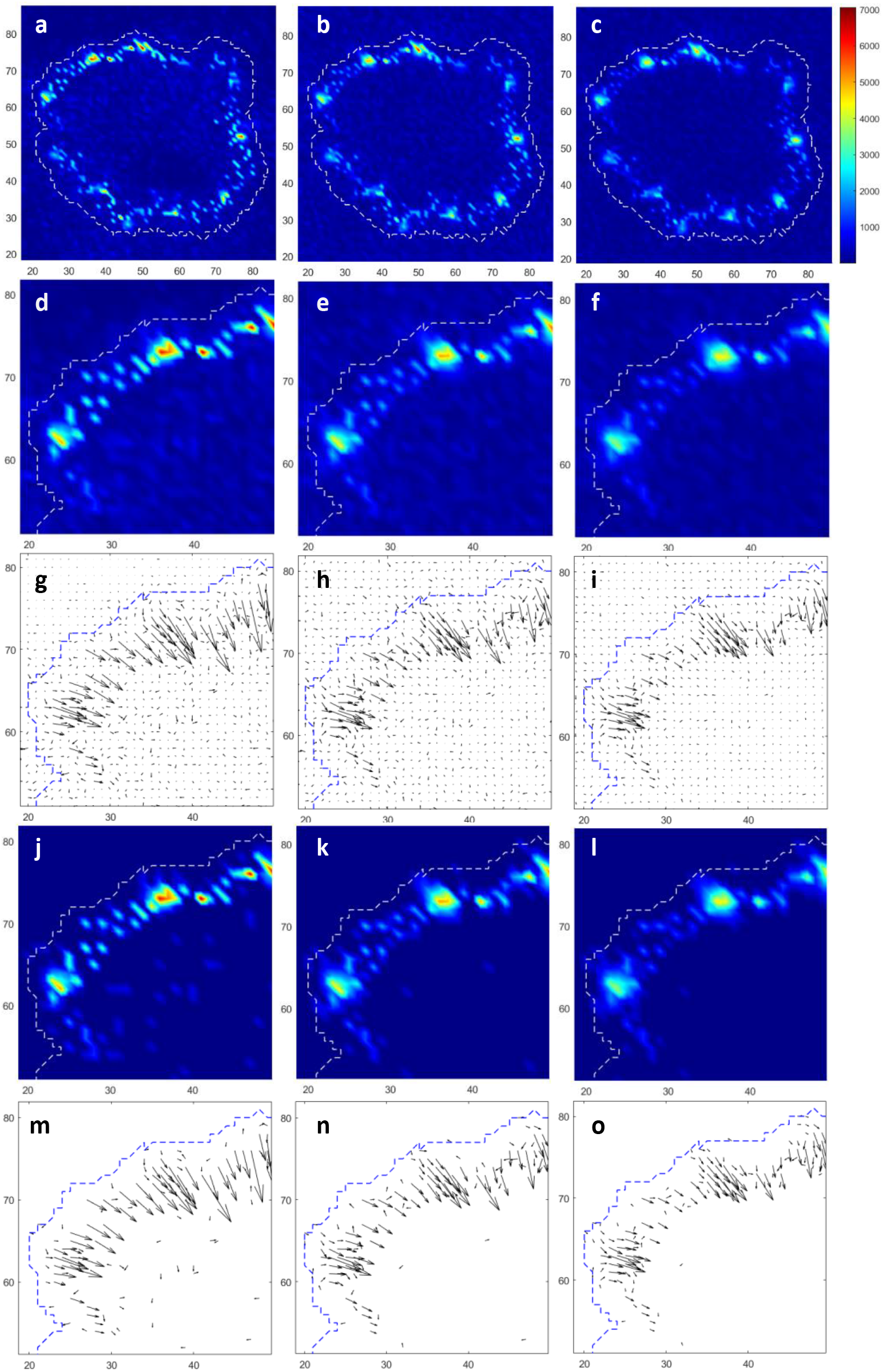
TFM of the same simulated testing cell as shown in Fig. 2, with a Gaussian noise of 0.0188 modeling pixels, equivalent to that of experimental measurements, added to the strain field. Stress fields are calculated with deep learning (left column), or FTTC with λ = 6×10^-10^ (middle column) or 1×10^-9^ (right column) for regularization, and are displayed as heat maps at a low (a-c) or high (d-f, j-l) magnification, or as vector plots (g-i, m-o), without (d-i) or with (j-o) cutoff filtering. Stress is calibrated as described in Fig. 2. The corresponding images of ground truth are shown in Fig. 2a, 2d and 2g.

The error for noisy data came at least in part from the background vectors that populated stress-free regions both inside and outside the cell (Fig. 4g, 4h, 4i). These vectors may be removed with a cutoff filter using the 98^th^ percentile stress outside the cell as the threshold (Fig. 4j, 4m), which resulted in a decrease in 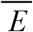 for deep learning TFM, by 43% and 64% respectively at 1x and 2x experimental noise respectively. With cutoff filtering, 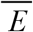 for deep learning TFM became smaller than 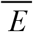 for FTTC regardless of regularization (Fig. 3, middle and right set, red textured bars; p < 0.001). Not surprisingly, the filter has the most prominent effect when noise was high and negligible effect with noise-free strains (Fig. 3, solid versus textured bars).

### Deep learning traction force microscopy for experimental data

We applied deep learning TFM to experimental strains measured with NIH 3T3 cells on polyacrylamide substrates. Heat maps showed a qualitative similarity between the stress field generated with deep learning TFM and FTTC without regularization (Fig. 5a-f). However, some structures appeared much more prominently in FTTC than in deep learning TFM (Fig. 5b, 5e, arrows). The application of regularization to FTTC at λ = 6×10^-10^ diminished these differences (Fig. 5b, 5h). However, more aggressive regularization at λ = 3×10^-9^ caused a serious loss of resolution (Fig. 5j-l).

**Figure 5.**
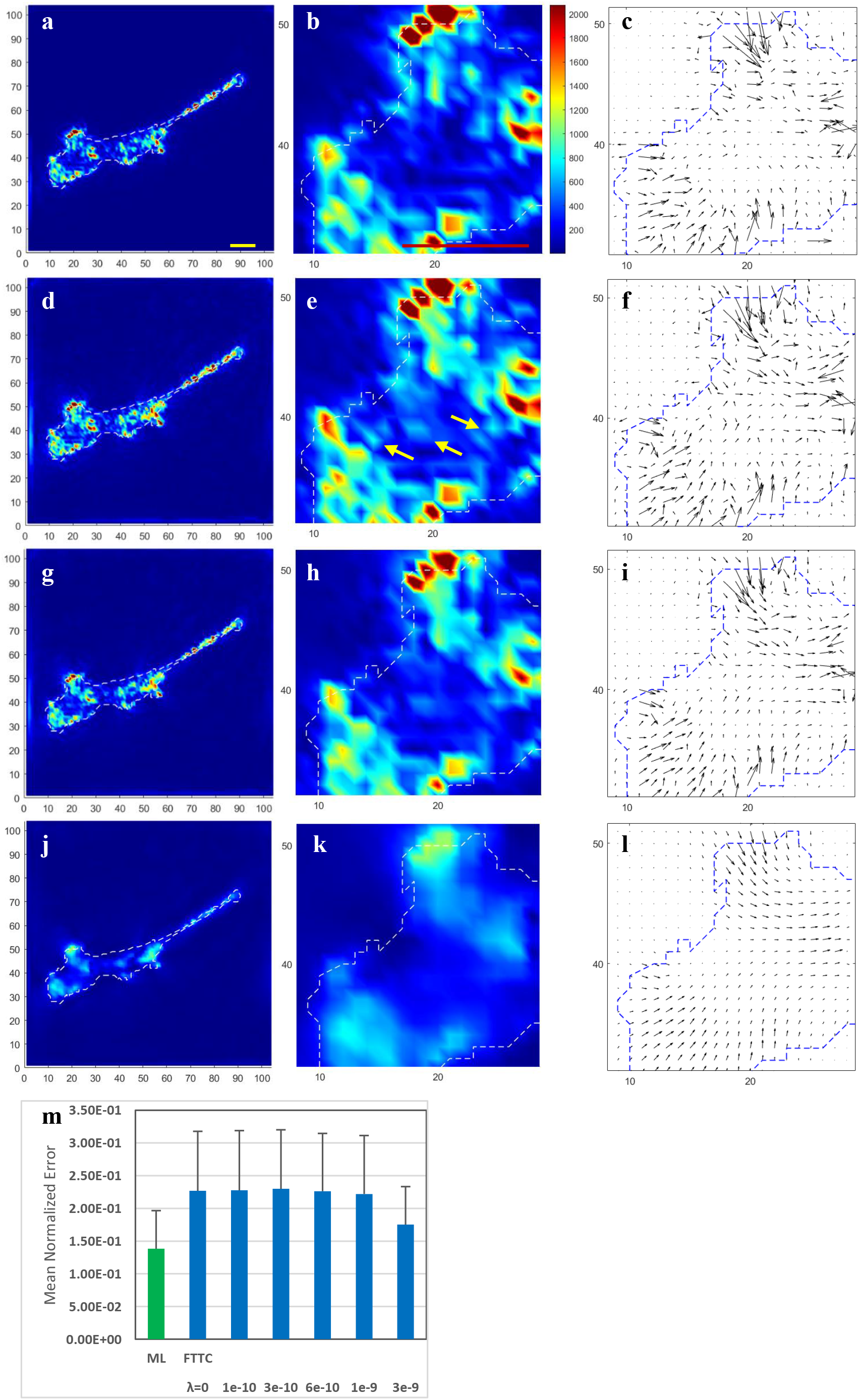
TFM of an NIH 3T3 cell plated on a polyacrylamide substrate of 10,670 Pa, embedded with fluorescent beads as strain markers. Stress fields are calculated with deep learning (a-c), or FTTC with λ = 0 (no regularization; d-f), λ = 6×10^-10^ (g-i), or λ = 3×10^-9^ (j-l), without applying cutoff filter. Heat maps of stress fields at a low magnification (a, d, g, j), or high magnification focusing on the frontal region (b, e, h, k), show a similar distribution of stress between deep learning TFM (a-c) and FTTC without regularization (d-f). However, some structures appeared much more prominently in FTTC than in deep learning TFM (e, arrows). Regularization at λ = 6 x10^-10^ diminishes these structures (d-i), while a high value of λ = 3×10^-9^ causes a serious loss of resolution (j-l). Mean normalized errors from 15 cells show a higher error for FTTC over a wide range of λ (m, blue bars) than deep learning (m, green bar). Error bars represent standard deviation. Scale bar, 20 μm (a, b) or 2 kPa (c, lower right).

Due to the lack of ground truth for experimental stress field, the accuracy of TFM was assessed quantitatively based on the deviation between measured strains and strains calculated from the calculated stress field, using an error equation similar to that for evaluating stress. The mean normalized error for deep learning TFM was 60% that for FTTC over a wide range of λ (Fig. 5m, p < 0.002), then decreased at λ = 3×10^-9^ (p < 0.03).

## Discussion

TFM belongs mathematically to an ill-posed inverse problem (17), where a unique solution may not exist and the solution is sensitive to noise. Conventional TFM relies on regularization to address this challenge (17), with the caveat that regularization may decrease the resolution and increases the deviation between measured strains and strains calculated from predicted stresses. In addition, it is difficult to define the optimal weight parameter λ for regularization (18).

Deep learning has emerged as an appealing approach for solving ill-posed inverse problems (19). Although deep learning also applies regularization, it was limited to the training process for the purpose of minimizing over-fitting (20), which suppresses the error for training at the expense of error for subsequent applications. When making predictions, deep learning works without regularization thereby avoiding the associated compromises. Interestingly, a higher accuracy with deep learning TFM than FTTC was also observed when processing noise-free strains without regularization. A possible cause is the difficulty in handling the singular central element of Green’s tensors in TFFC (14), while the effect of varying its value appeared small for deep learning.

The application of deep learning hinges on two major requirements, a network architecture compatible with the problem, and the availability of a sizeable training dataset. We have adapted UNet for processing 2D fields of stress and strain vectors (22), using different z planes in a 3D matrix to represent different orthogonal vector components. In addition, since the strain field along a given direction is determined by stresses along both orthogonal directions, it is essential to replace 2D convolution with 3D convolution in order to generate the required linear combinations.

For the training dataset, we have adapted a simulation model for cell migration to generate unlimited pairs of stress and strain fields (21), under the assumptions that all stress vectors point toward the cell center and have a magnitude proportional to the net signal that determines protrusion or retraction activities. While these assumptions may represent oversimplifications, they allowed the generation of simulated fields that resembled cellular stress fields for successful training, as evidenced by the similar predictions of deep learning TFM and FTTC where the fundamentally different mathematical approaches argued for the validity of both methods.

Simulated data, with their known ground truth stress fields, further allowed rigorous analysis of the performance of deep learning TFM under defined noise conditions. We observed a small mean normalized error of 3.2% for stress fields generated from noise-free strains. In addition, all the qualitative features were preserved even when the noise of strains was twice that of experimental measurements. A visible effect of noise was the generation of small background stress throughout the image, which may be removed by applying a cutoff filter without affecting the resolution or structural features. Similar background also appeared in stress fields generated with FTTC, which may be suppressed either by regularization, with a loss of resolution and features, or using a similar cutoff filter. With cutoff filtering, deep learning TFM showed a better accuracy than FTTC under all the tested conditions.

While other conventional TFM methods may yield a better accuracy and/or resolution than FTTC as tested in the present study (25, 26), the speed of deep learning TFM at < 1 ms for 104×104 modeling pixels, is comparable to that of the fastest conventional TFM methods. In addition, a neural network pre-trained for a given Poisson ratio may be scaled easily for different Young’s moduli and magnifications. Additional advantages of deep learning include its ability to generate stress fields without prior knowledge of the location of cell border or focal adhesions.

While deep learning does not enforce the balance of total forces, the small normalized error as demonstrated with simulated testing data argues for its general reliability. In summary, we showed that deep learning TFM can serve as an appealing alternative to conventional 2D TFM for characterizing cell-substrate mechanical interactions at a high speed, resolution, and accuracy.

## Author Contributions

Yu-li Wang performed all the computation work, including the generation of simulated data, the processing and analysis of experimental data, and the development and training of neural network. Yun-chu Lin performed cell culture, prepared elastic substrates, and collected the experimental strain data.

## Acknowledgements

We thank Mr. Xinhan Li for drafting the neural network architecture. This study was supported by NIH grant R01 GM118998.

## Supplemental Information

**Supplemental Figure 1.**
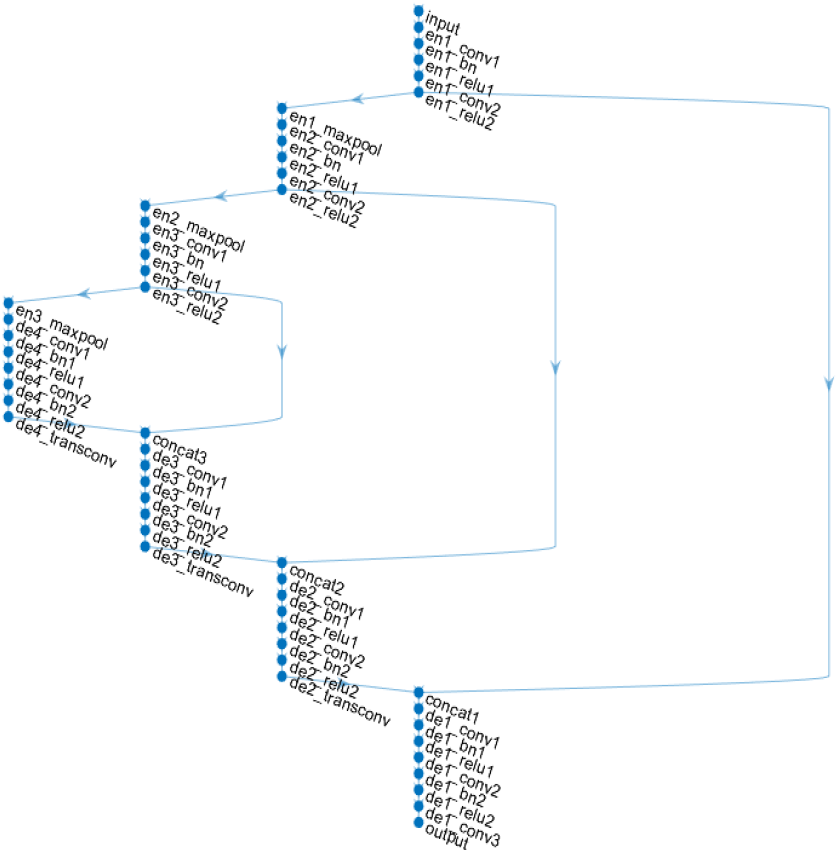
Modified UNet neural network for deep learning TFM, showing 51 layers with ‘enN’ denoting encoding levels and ‘deN’ decoding levels where N is level number. Each level contains two 3D convolution layers (denoted as convN), except for the bottom level where there is an additional convolution layer before the output regression layer. Each decoding level ends with a layer of transposed 3D convolution (denoted as transconv) for up-sampling. Other layers perform batch normalization (denoted as bnN), rectified linear unit operation (denoted as reluN), and max pooling (denoted as maxpool), where N is level number. Three skip connections are indicated as vertical shunts that concatenate matrices of compatible dimensions from encoding and decoding levels, for the purpose of preserving the resolution. The neural network exits with a regression output layer.

**Supplemental Figure 2.**
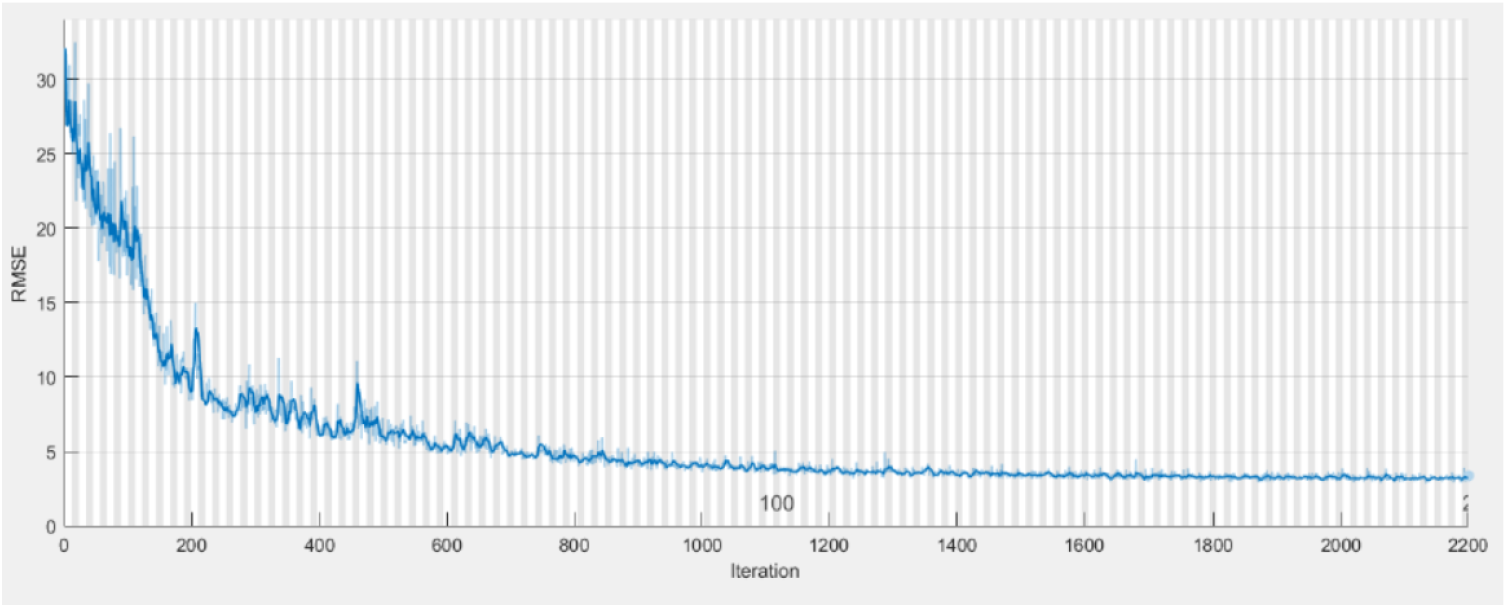
Typical training progress of the neural network for deep learning TFM. The training dataset consists of 708 images, divided into minibatches of 64 images each with random shuffling at each epoch. Thus, each epoch (upper number along the x-axis) consists of 11 minibatch iterations (lower number along the x-axis). Y-axis shows the root mean squared error, calculated as the square of error at each modeling pixel summed over the entire area of 104×104 modeling pixels before taking the square root and averaged over the images in the minibatch.

